# Cation permeability and pore dynamics in TRPV1 ion channels

**DOI:** 10.1101/2023.05.25.542342

**Authors:** Miriam García-Ávila, Javier Tello-Marmolejo, Tamara Rosenbaum, León D. Islas

## Abstract

The Transient Receptor Vanilloid 1 (TRPV1) is a non-selective ion channel, which is activated by several chemical ligands and heat. We have previously shown that activation of TRPV1 by different ligands result in single-channel openings with different conductance, suggesting that the selectivity filter is highly dynamic. TRPV1 is weakly voltage-dependent, here we sought to explore whether the permeation of different monovalent ions could influence the voltage-dependence of this ion channel. By using single-channel recordings, we show that TRPV1 channels undergo rapid transitions to closed states that are directly connected to the open state, which may result from structural fluctuations of their selectivity filters. Moreover, we demonstrate that the rates of these transitions are strongly influenced by the permeant ion, suggesting that ion permeation regulates the voltage dependence of these channels.

## Introduction

Transient receptor potential vanilloid (TRPV) channels are members of a superfamily of voltage-gated-like ion channels (Yu and Catterall, 2004). The first member of the TRPV family to be identified was TRPV1, which is a non-selective cationic channel involved in thermal sensation and which participates in the physiology of pain and inflammation (Rosenbaum et al., 2022). TRPV1 channels assemble as homotetramers, and each subunit contains large intracellular N-and C-terminal domains of complex architecture (Liao et al., 2013), and a transmembrane domain formed by six alpha helical membrane-spanning segments. Within this transmembrane domain, the transmembrane helices S1 to S4 form a helix bundle constituting a voltage sensor-like domain (VSLD), that does not seem to function as the voltage-sensor domains (VSD) present in canonical voltage-activated channels (Palovcak et al., 2015a), but which contains multiple binding sites for activating and regulatory molecules (Cao, 2020). The pore, which <u>encompasses</u> a selectivity filter and an activation gate, is formed by the S5 segment, a pore loop and a helix motif and the S6 segment (Liao et al., 2013).

TRPV1 can be activated by varied stimuli such as capsaicin, external pH < 5.4, heat, various ligands and voltage (Caterina and Julius, 2001; Jordt et al., 2000; Matta and Ahern, 2007; Grandl et al., 2010; Rosenbaum et al., 2022). Interactions with these stimuli lead to widening of the S6 activation gate, culminating in channel opening and ion permeation (Salazar et al., 2009). Importantly, these activating factors are allosterically coupled to the channel gate and also between them (Jara-Oseguera and Swartz, 2015).

Several structures of TRPV1 and related channels (TRPV1-4) have been solved by X-ray crystallography and cryoEM methods (Liao et al., 2013; Zubcevic et al., 2018; Shimada et al., 2020; Deng et al., 2018). These structures show that in the presence of activators, the selectivity filter (SF) can adopt varying conformations, suggesting that in these channels the SF is a highly dynamic structure (Zubcevic et al., 2018). Functional experiments have shown that, while not an activation gate, the SF shows dynamic accessibility changes (Jara-Oseguera et al., 2019). It has also been suggested that voltage-induced conformational changes in this structure underlie the very small voltage dependence of activation (Yang et al., 2020).

The dynamic character of the selectivity filter of TRPV1 has not been studied functionally, although several lines of evidence point towards this characteristic. For example, different agonists of TRPV1 can open the channel to distinct single-channel conductance levels (Canul-Sánchez et al., 2018; Geron et al., 2018). <u>Hence, we</u> decided to characterize the effect of the permeant ions on the gating of TRPV1. We observed very rapid transitions to closed states that are directly connected to the open state and that might represent structural fluctuations of the SF. The rates of these transitions are strongly influenced by the permeant ion, in a manner that is consistent with ion transport being involved in the regulation of gating of these channels.

## Methods

### Cell culture and transfection

We used the HEK293 cell line from American Type Culture Collection (ATCC; RRID: CVCL_0045) for heterologous expression of TRPV1. Cells were grown in Dulbecco’s Modified Eagle Medium (DMEM, Invitrogen) with the addition of 10 % fetal bovine serum (Invitrogen) and 1% of penicillin-streptomycin (Invitrogen) (referred to as supplemented DMEM). Cultures were incubated at 37°C, in a 5 % CO_2_ atmosphere. Every 3 or 4 days, the cells were washed with PBS, followed by treatment with 1 ml of 0.05% trypsin-EDTA (Invitrogen) for 2 min, then 1 ml of supplemented DMEM was added. Subsequently, the cells were mechanically dislodged and 80 μl of the cell suspension were reseeded on 5 x 5 mm glass coverslips in 30 mm culture dishes with 2 ml of supplemented DMEM. After 1-2 days in culture, cells were cotransfected with pcDNA3.1 plasmid with WT rat TRPV1 (rTRPV1) and pEGFP-N1 plasmid containing the eGFP fluorescent protein using the jetPEI transfection reagent (Polyplus), according to manufacturer’s instructions. Single-channel recordings were done one day after transfection, while macroscopic recordings were performed two or more days after transfection.

### Electrophysiological recordings

The experiments were carried out in the inside-out configuration of the patch-clamp recording technique (Hamill et al., 1981). Recordings were performed using a pipette (extracellular) solution consisting of 130 mM NaCl, 10 mM HEPES and 5 mM EGTA, pH 7.4, and bath (intracellular) solutions with either 130 mM NaCl, KCl, RbCl, LiCl, NH_4_Cl or CsCl (depending on the experiment, as indicated in the figure legends), 10 mM HEPES, and 5 mM EGTA, pH 7.4.

Currents were recorded with an EPC-10 patch-clamp amplifier (RRID:SCR_018399, HEKA Elektronik) controlled by PatchMaster software (RRID:SCR_000034, HEKA Elektronik). Borosilicate capillaries (Sutter instruments) were used to fabricate patch-pipettes, which had a resistance after fire polishing of 4-6 MΩ for macroscopic currents and 8-10 MΩ for single-channel recordings.

Macroscopic currents were filtered <u>with the build-in 4-pole Bessel filter of the EPC-10</u> at 1 kHz <u>(−3dB) and sampled at 40 kHz while single-channel currents were filtered at 5 kHz and sampled at 50 kHz.</u>

### Analysis of macroscopic current recordings

The macroscopic conductance in each permeant ion condition was determined from the current *I*, applying Ohm’s law according to:

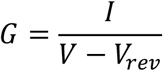

where V is the test voltage and V_rev_ is the reversal potential. The conductance was normalized to the maximum G_max_ and plotted as a function of voltage and was fit to a Boltzmann function:

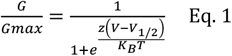

where *z* is the apparent valence of gating in e_o_, V_1/2_ is the potential where *G/G*_*max*_=0.5, *K*_*B*_ is Boltzmann’s constant and *T* the temperature in Kelvin (296 K). <u>The apparent free energy change of activation by voltage was estimated as ΔG_ion_ = zV_1/2_ from values obtained for each ion from fits to Eq. 1. ΔΔG was estimated as ΔG_ion_-ΔG_Na_</u>

### Analysis of single-channel recordings

Single-channel recordings were analyzed to determine the open probability, P_o_. Openings were detected employing the 50% threshold crossing method (Colquhoun and Sigworth, 1995a). Single-channel traces were interpolated with a cubic spline function before event detection. Dwell times (d_o_) of open events and dwell times of closed events (d_c_) were compiled and used to determine *P*_*o*_ as:

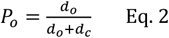

The dwell times were also used to construct amplitude vs. open dwell time relationships.

<u>Single-channel kinetics were analyzed by constructing open and closed dwell time distributions using log-transformed histograms according to (Sigworth and Sine, 1987). Events shorter than the filter death time T_d_ were discarded. T_d_ = 0.179/f_c_. At f_c_ = 5 kHz, T_d_ = 36 μs. Bursts were defined as groups of openings separated by a closed time longer than a critical time, t_c_. This critical time was calculated from the closed time distribution from recordings in Li^+^, Na^+^, Rb^+^ and Cs^+^ according to (Colquhoun and Sakmann, 1985). Closed, open and burst length distributions were fit to a sum of exponential components. The number of components and their parameters are given in the figure legends.</u>

### Analysis of fast single-channel current fluctuations

Fast current fluctuations are apparent in single-channel records both as increases in the open-channel noise and as incomplete closing transitions. Since the half-amplitude crossing method is not appropriate to study these extremely fast events, we chose to analyze these fast kinetics by studying the all-points amplitude histograms compiled from bursts of channel activity, employing an extension of the Beta-distribution analysis developed by (Schroeder, 2015), which <u>has been shown to be</u> applicable to systems with more than two states. This analysis relies on fitting of the all-points amplitude histograms to theoretical distributions derived from time series of simulated channel activity from specific kinetic models. These fits yield forward and backward rate constants that were analyzed over varying conditions of permeant ion and voltages. <u>Bursts of channel openings were selected manually from recordings and included short periods of no channel activity in order to adjust the zero current level. Experimental all-points histograms were constructed by accumulating the currents from these selected bursts.</u>

### Model fitting

A simulated time series of channel openings and closings from specific Markov models of gating was first generated employing a Monte Carlo method. For exponentially distributed dwell times, any particular random duration of value *d*_*i*_, in any closed or open state, *i*, is calculated from rate constants as:

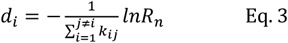

Here, *k*_*ij*_ are the rate constants leaving state *i* into state *j* and *R*_*n*_ is a uniformly distributed random number from 0 to 1. Transition probabilities between sates *i* and *j, p*_*ij*_ were calculated according to:

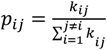

Once a time series is simulated, <u>Gaussian noise with a variance similar to the variance of the closed channel current level of each recording was added and the time series was</u> filtered by convolution with the impulse response of a Gaussian filter given by (Colquhoun and Sigworth, 1995b):

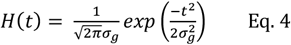

σ_g_ is related to the filter corner frequency *f*_*c*_ by:

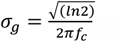

Simulated and filtered data are converted to an all-points amplitude histogram which is then fitted to the experimental histogram using a least-squares procedure and the fit parameters used to simulate a new time series until the fit error is minimized. <u>We implemented a zero-order optimization algorithm with a number of iterations i = 100. The merit function was R_n_(i+1) < R_n_(i), where R_n_(i) is an average of n least squares values between a simulation and the experimental data. The number of simulations per iteration, was n = 10. It was essential to average these least squares values R_n_(i) since the simulation is a random process and non-optimal values could produce a low R because of this randomness. Two types of steps were taken with equal probability in the variation of the fitting parameters. A small step that looks for the minimum R value and a large step. The large step was important to avoid getting stuck in local minima.</u>

Experimental amplitude histograms were fitted to simulated histograms resulting from several models considering one open estate and one, two or more closed states with appropriate voltage dependent rate constants.

### Fit to rate constants

The voltage-dependence of each transition was calculated by plotting the rate constants as a function of voltage and fitting the data to the following function:

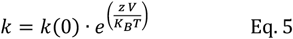

Here, *k*(0) is the rate constant at 0 mV, z is the apparent charge associated with the transition, *K*_*B*_ is the Boltzmann constant, *T* is the temperature (296 K) and *V* is the voltage applied.

#### Fast transitions showed saturation at high voltages and were thus fitted using a subtraction of two exponential functions

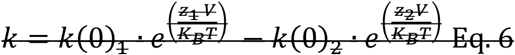

Data were analyzed with Patchmaster (HEKA) and custom-written programs in Igor Pro 8 (RRID: SCR_000325, Wave-Metrics) and Python 3 using Jupyter Notebooks (RRID:SCR_008394).

### Statistical tests

Statistical comparisons were made by using one-way ANOVA with significance determined from Tukey’s post hoc test. *p < 0.05 indicates statistical significance.

## Results

In these experiments, we focused on outward currents carried by varying permeant ions, in the nominal absence of divalent ions and presence of a constant extracellular concentration of NaCl. These conditions were chosen since it has been shown that Ca^2+^ can permeate, block and produce Ca^2+^-calmodulin mediated desensitization of TRPV1 (Rosenbaum et al., 2004; Numazaki et al., 2003) and that extracellular Na^+^ ions are required for normal gating in TRPV1 (Jara-Oseguera et al., 2016).

### Permeation and voltage dependence of TRPV1

Macroscopic currents through TRPV1 were activated by a saturating concentration of capsaicin (CAP) of 10 μM and recorded in asymmetric ion concentrations, with Na^+^ always present at 130 mM in the extracellular face of the channel. Outward currents were carried by the test ion and were always preceded by recording of current in symmetric Na^+^ in the same patch, allowing the results to be compared over multiple patches, taking as reference the condition in which the channel permeates Na^+^ in both directions. For example, red traces for only Na^+^ vs. blue traces for Na^+^/Li^+^ (Figure 1A). As can be seen from I-V curves normalized to the current in Na^+^ at 130 mV, TRPV1 permeates monovalent ions (Figure 1B) (Caterina et al., 1997a).

**Figure 1.**
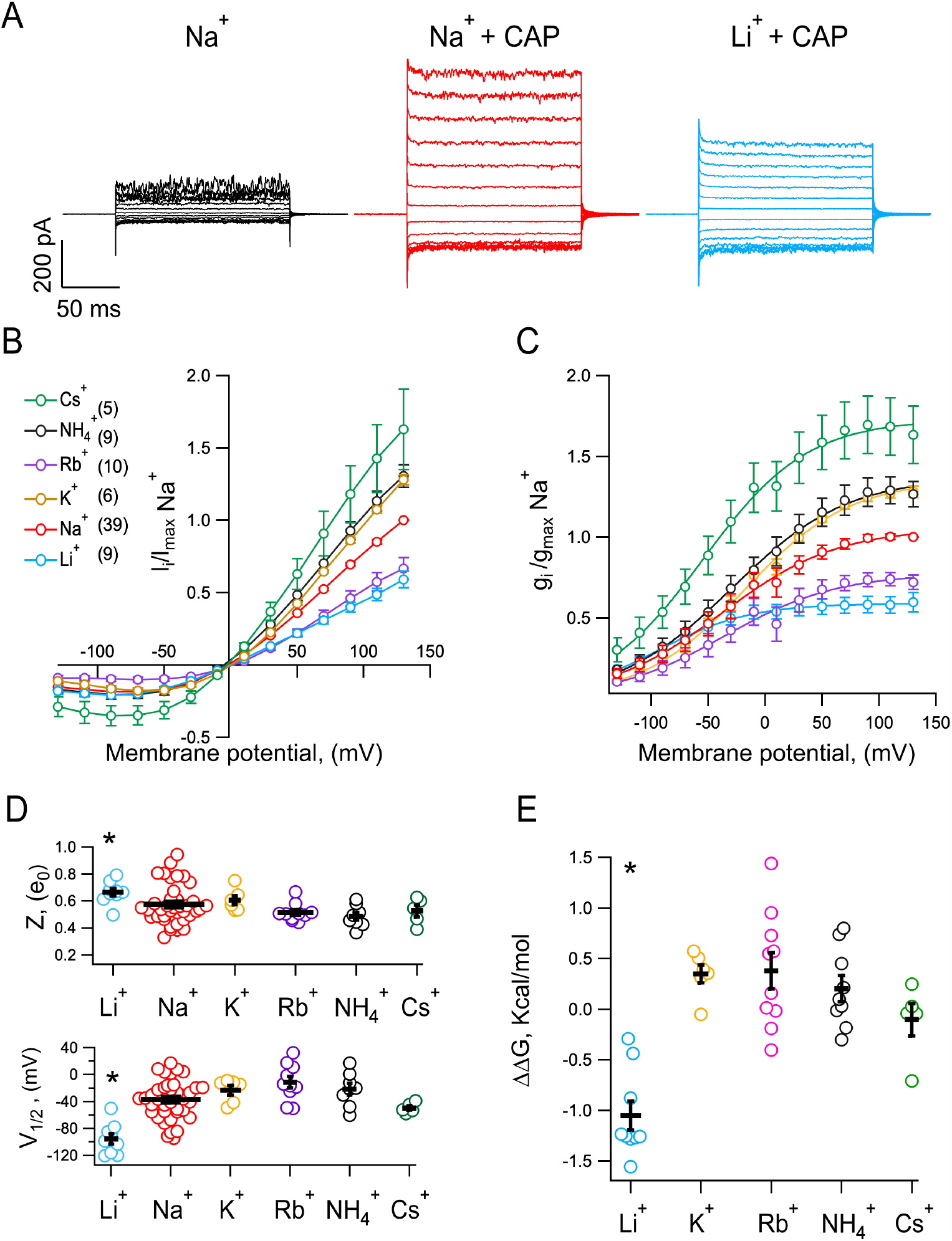
The voltage-dependence of TRPV1 is influenced by the permeant ion. **A)** Representative inside-out current-traces recorded in the same patch, elicited by 20 mV steps from −130 mV to 130 mV for a duration of 150 ms. The extracellular solution (pipette) contained 130 mM NaCl, this patch was first exposed to internal 130 mM NaCl (black) followed by exposure to 130 mM NaCl + 10 μM capsaicin (red) and finally to 130 mM LiCl + 10 μM capsaicin (blue). All experiments were performed following the same experimental strategy to compare the permeation of different monovalent ions (Li^+^, K^+^, Rb^+^, NH_4_^+^, Cs^+^) with respect to the permeation of Na^+^ in the same patch. **B)** Averages of current-voltage (I-V) relations for all ions tested. The number of experiments, n, is in parenthesis and is the same for panels C-E. **C)** Conductance-voltage (G-V) relations for monovalent ions, obtained from data as in B in the presence of 10 μM capsaicin, normalized to data with a symmetrical concentration of Na^+^ from each patch. G-V relations were fitted to equation 1 (continuous lines). **D)** Scatter plots of the apparent charge *z* (above) and V_½_ (below) associated with channel opening, both were estimated by fitting G-V curves to equation 1. The values of *z* for Li^+^ were compared for statistical differences with all other ions, the p values are: *p = 0.00007 for Li^+^ vs. Na^+^; NS (not significant) p=0.213 for Li^+^ vs. K^+^; p = 0.00034 for Li^+^ vs. Rb^+^; p = 0.00005 for Li^+^ vs. 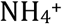; P = 0.01 for Li^+^ vs. Cs^+^. For V_½_: p = 0.0000009 for Li^+^ vs. Na^+^; p = 0.00001 for Li^+^ vs. K^+^; p = 0.00000001 for Li^+^ vs. Rb^+^; P = 0.000002 for Li^+^ vs. 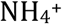; p = 0.02 for Li^+^ vs. Cs^+^. **E)** Scatter plot of the ΔΔ*G* associated to the ion effect in channel opening was estimated as: 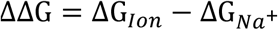. The * symbol indicates statistical differences, p = 0.00005 for Li^+^ vs. K^+^; p = 0.0000002 for Li^+^ vs. Rb^+^; p = 0.000004 for Li^+^ vs. 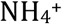; p = 0.003 for Li^+^ vs. Cs^+^. Data are shown as mean ± SEM. The significant difference was determined with a one-way ANOVA test followed by a Tukey’s test post-hoc.

Outward macroscopic currents are larger when the permeant ion has a larger ionic radius like Cs^+^ (1.6 Å). However, the reversal potential was not statistically different from zero for each ion tested (Figure 1B), showing that the channel does not select between monovalent ions. Conductance (G) vs. voltage (V) relationships, again, normalized to G_Na_, show that the magnitude of the maximal conductance at positive voltages (G_max_) depends on the permeant ion. These G-V curves were fitted to Eq. 1, to estimate apparent charge, z and voltage of half activation V_1/2_. Besides the effect on G_max_, the values of z and V_1/2_, are different for each of the permeant ions studied; specifically, there is a significant difference in the ΔΔG with Li^+^ (Compared to Na^+^, Figure 1, D and E), which is brought about by a significant increase in z (increased steepness of the G-V curve) and a negative shift of the V_1/2_.

To understand why the macroscopic conductance is affected by the permeant ion, we performed single-channel recordings in identical ionic conditions. Our single-channel recordings in the inside-out configuration show a clear increase in the microscopic conductance when larger ions permeate (Figure 2A). Since the reversal potential was very close to zero for all ions (Figure 2B), to estimate the cation permeability, we used the magnitude of the single-channel conductance, relative to the single-channel conductance of Na^+^ current in the same patch. We can estimate the permeability sequence as: Li^+^ < Na^+^ < K^+^ < Rb^+^ < 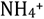 < Cs^+^ (Figure 2C).

**Figure 2.**
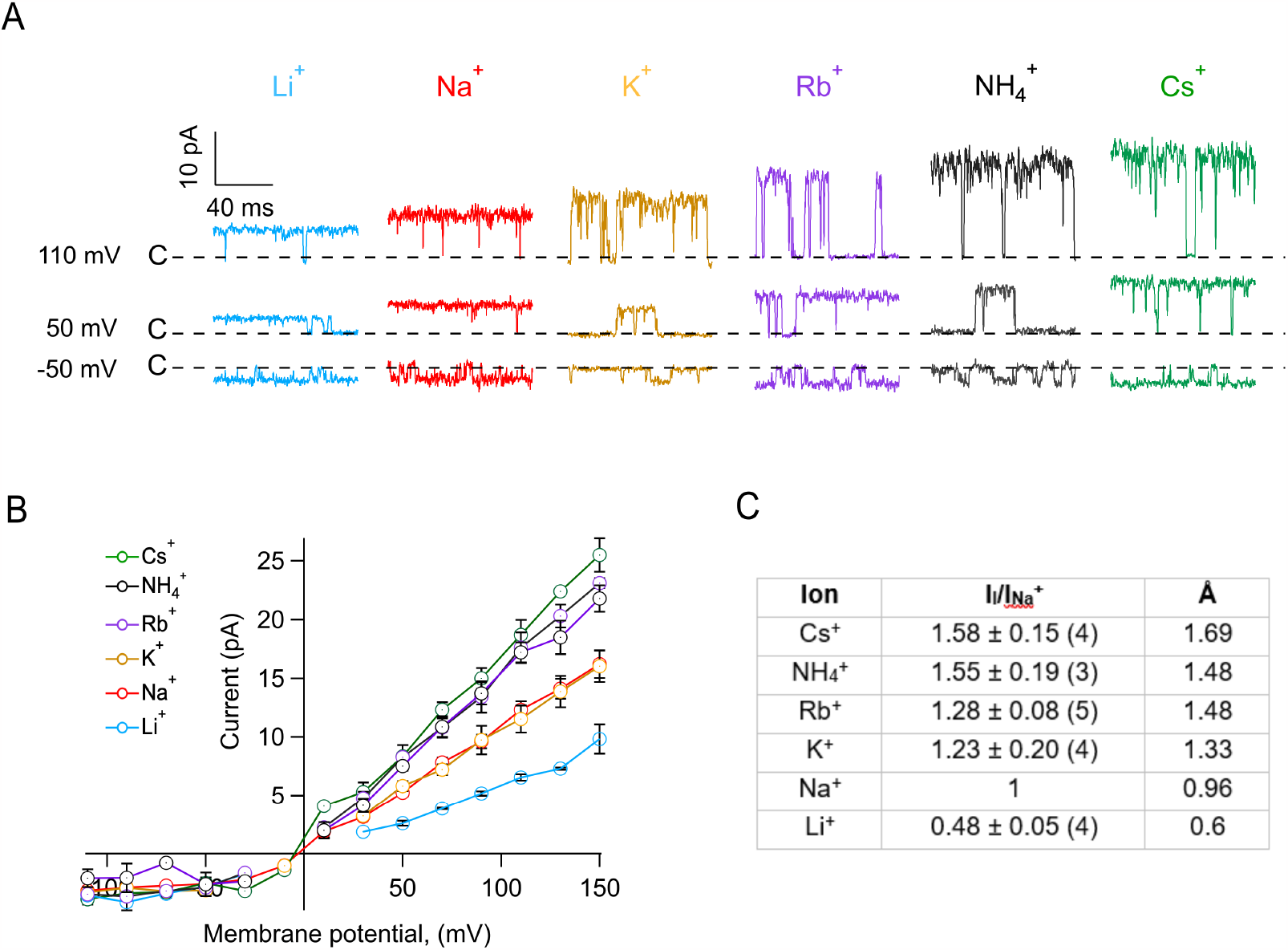
TRPV1 permeability to monovalent ions is size-dependent. **A)** Representative current traces showing openings from single-channel recordings in the presence of varying permeant ions. Inside-out patches were first exposed to internal 130 mM Na^+^ + 100 nM capsaicin and then to internal 130 mM (Li^+^ (blue), Na^+^ (red) K^+^(mustard), Rb^+^ (purple), 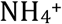(black), Cs^+^ (green)) + 100 nM capsaicin, similarly to experiments in Figure 1A. The dashed line represents the closed state of the channel.<u>Currents are shown filtered at 2.5 kHz, for clarity of illustration.</u> **B)** I-V relations from single-channel records. These were obtained from data as in A for each ion. **C)** Current carried by each ion relative to the single-channel current amplitude with Na^+^. All currents measured at 90 mV. Ionic radii in A’ are also shown for comparison. Data are mean ± s.e.m., n is shown in parentheses and is the same for the curves in **B**.

Although the single-channel conductance explains in part the larger macroscopic conductance when the permeant ion has a larger atomic radius, i.e., larger single-channel conductance corresponds to larger macroscopic conductance, the microscopic current-amplitudes observed by measuring single-channels (Figure 2B) do not fully correspond to what is observed in macroscopic currents (Figure 1C). Specifically, the single-channel conductance with Rb^+^ is greater than in K^+^, Na^+^ and Li^+^, unlike the macroscopic results, where the conductance with Rb^+^ is almost the lowest, only comparable with the conductance with Li^+^.

This difference suggests that when Rb^+^ is the permeant ion at positive voltages, the channel might gate differently. To test this possibility, we measured the single-channel open probability according to Eq. 2, with different permeant ions. Figure 3A shows that in the presence of Rb^+^, the channel opens in short bursts, unlike the openings in Na^+^, which happen in long-lived bursts <u>at this concentration of capsaicin</u>. Concurrently, the open probability is significantly smaller with Rb^+^, while it is close to the maximum <u>attained P_o_ at this capsaicin concentration (100 nM) and voltage in all other ions tested (Figure 3B). Since the macroscopic current is proportional to the product of P_o_ and single channel I-V,</u> this result reconciles the observed effects of ions on microscopic and macroscopic conductance and reveals that the permeant ion modulates gating of TRPV1. In particular, Rb^+^ makes TRPV1 gate with lower open probability.

**Figure 3.**
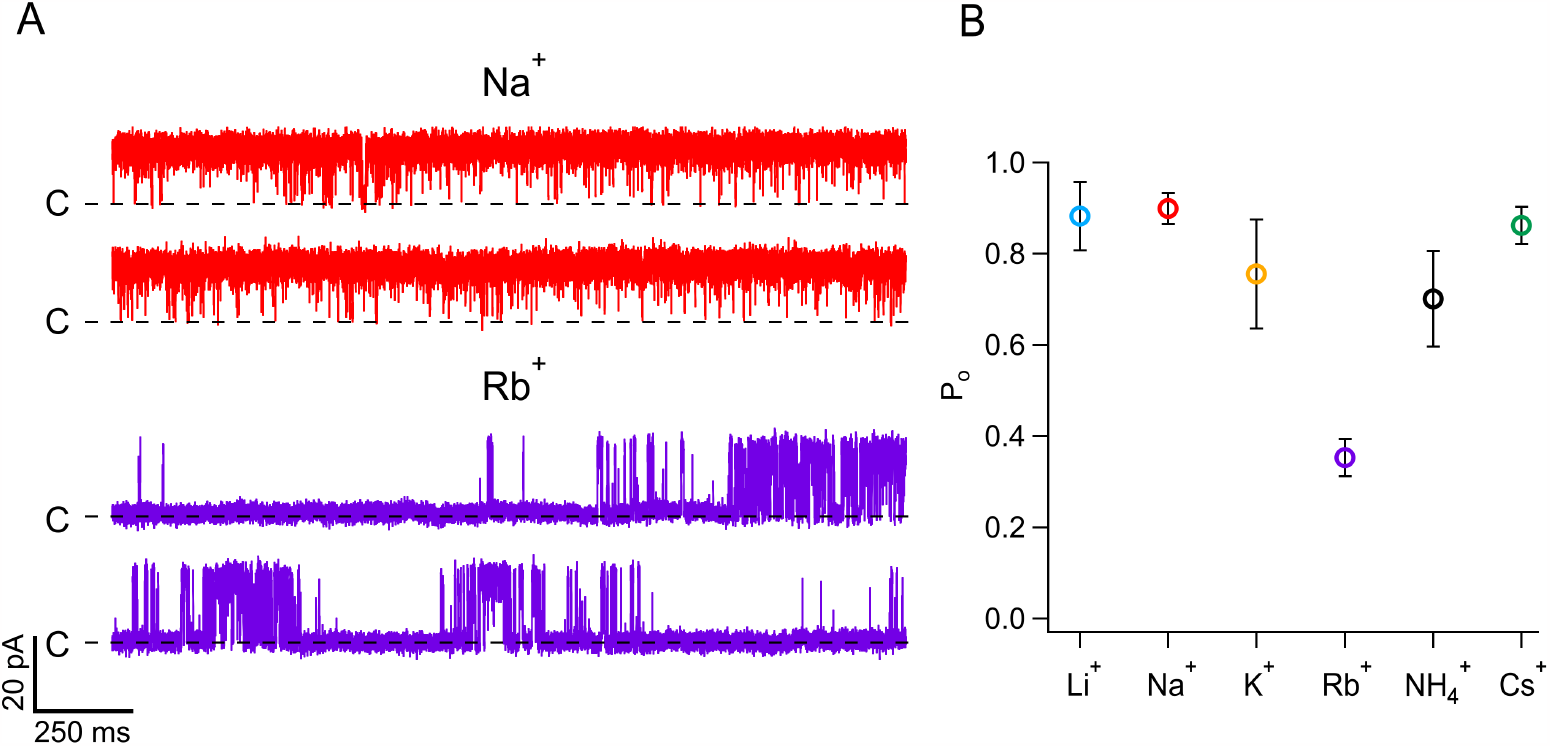
Rubidium ions strongly modulate the single-channel open probability of TRPV1 channels. **A)** Representative current traces from single-channel recordings at 70 mV when intracellular Rb^+^ (purple) or Na^+^ (red) were used as permeant ions. <u>Currents are shown at full bandwidth (5 kHz).</u> **B)** Open probability of TRPV1 channels at <u>70 mV</u> in the presence of different permeant ions and 100 nM of capsaicin, calculated from traces of current recordings as shown in A. Data are mean ± s.e.m. Number of experiments is: Li^+^(4); Na^+^(8); K^+^(3); Rb^+^(5); 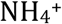(4); Cs^+^(4).

<u>To further explore the gating effects of permeant ions, we performed an analysis of the kinetics of channel opening and closing. TRPV1 gating at the single-channel level is complex (Hui et al., 2003), the open and closed dwell time histograms can be fitted by a sum of three and four exponential components, respectively, indicating the presence of multiple open and closed states (Supplementary figure 1 and Table I), with the channel mostly opening in bursts. For comparison, we obtained the burst length</u> distributions in Li^+^, Na^+^, Cs^+^ and Rb^+^. As shown in Figure 4, burst length is also multiexponential and the bursts become shorter when Rb^+^ is the permeant ion. Burst length distributions in Na^+^, Li^+^ and Cs^+^ are very similar.

**Table 1.** Time constants and amplitudes of individual exponential components fitted to the histograms in supplementary figure 1.

| Permeant ion | Open time |  | Closed time |  |
| --- | --- | --- | --- | --- |
|  | Time constant (s) | Amplitude | Time constant (s) | Amplitude |
| Li<sup>+</sup> | $\tau_1 = 0.0002$<br>$\tau_2 = 0.004$<br>$\tau_3 = 0.02$ | $a_1 = 2.3$<br>$a_2 = 3.4$<br>$a_3 = 4.8$ | $\tau_1 = 0.0001$<br>$\tau_2 = 0.0005$<br>$\tau_3 = 0.005$<br>$\tau_4 = 0.06$ | $a_1 = 7.5$<br>$a_2 = 4.2$<br>$a_3 = 2.5$<br>$a_4 = 1.1$ |
| Na<sup>+</sup> | $\tau_1 = 0.0001$<br>$\tau_2 = 0.0015$<br>$\tau_3 = 0.011$ | $a_1 = 11.6$<br>$a_2 = 2.3$<br>$a_3 = 16.6$ | $\tau_1 = 6.7 \times 10^{-5}$<br>$\tau_2 = 0.0005$<br>$\tau_3 = 0.0033$<br>$\tau_4 = 0.0305$ | $a_1 = 21.4$<br>$a_2 = 11.8$<br>$a_3 = 3.7$<br>$a_4 = 0.02$ |
| Rb<sup>+</sup> | $\tau_1 = 8.71 \times 10^{-5}$<br>$\tau_2 = 0.0011$<br>$\tau_3 = 0.003$ | $a_1 = 13.1$<br>$a_2 = 12.5$<br>$a_3 = 21.5$ | $\tau_1 = 6.6 \times 10^{-5}$<br>$\tau_2 = 0.0007$<br>$\tau_3 = 0.007$<br>$\tau_4 = 0.21$ | $a_1 = 27.1$<br>$a_2 = 15.9$<br>$a_3 = 10.2$<br>$a_4 = 4.1$ |
| Cs<sup>+</sup> | $\tau_1 = 0.0001$<br>$\tau_2 = 0.0071$<br>$\tau_3 = 0.0211$ | $a_1 = 16.5$<br>$a_2 = 25$<br>$a_3 = 5.6$ | $\tau_1 = 8.5 \times 10^{-5}$<br>$\tau_2 = 0.0009$<br>$\tau_3 = 0.0038$<br>$\tau_4 = 0.023$ | $a_1 = 31.5$<br>$a_2 = 15.5$<br>$a_3 = 6.2$<br>$a_4 = 2$ |

**Figure 4.**
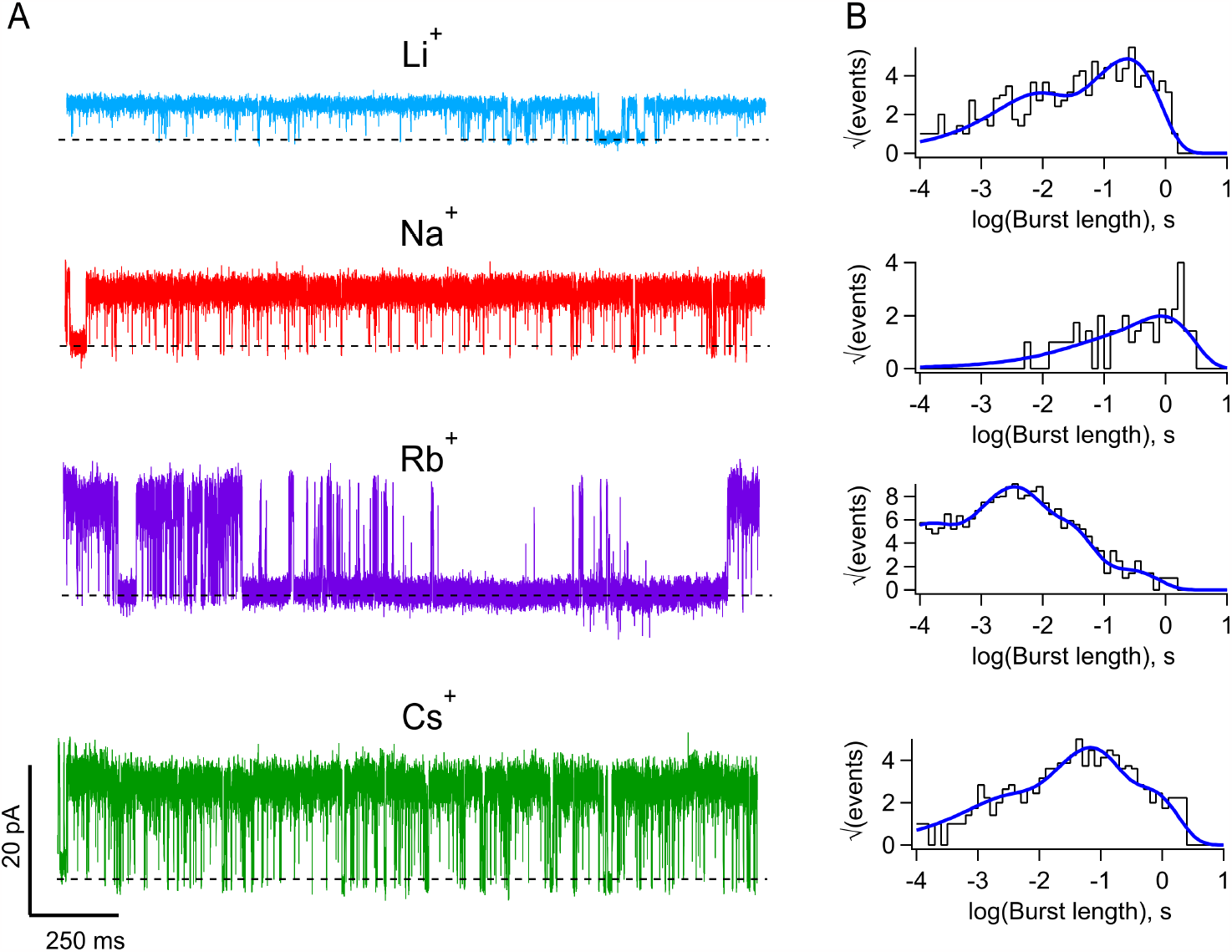
The permeant ion modulates single channel kinetics of TRPV1 channels. **A)** Representative single-channel traces with the indicated permeant ion. Data is shown at full bandwidth (5 kHz) except Li^+^ currents, which are filtered at 2.5 <u>kHz.</u> **B)** Burst length distributions corresponding to permeant ions in A. The continuous curve is the sum of exponential components. The values of time constants (τ, in s) and amplitudes (a_i_) are: Cs^+^; a_1_=1.5474, τ_1_ = 0.0013, a_2_=5.2727, τ_2_=0.04818, a_3_=4.3972, τ_3_=0.5031. Na^+^; a_1_=0.3720, τ_1_=0.0362, a_2_=3.2643, τ_2_=0.8346. Li^+^; a_1_=2.9447,τ_1_=0.0048, a_2_=8.0241,τ_2_=0.2362. Rb^+^; a_1_=5.0377, τ_1_=7.244 x 10^−5^, a_2_=8.7838,τ_2_=0.0021, a_3_=8.4035,τ_3_=0.0163, a_4_=2.8719, τ_4_=0.2094.

TRPV1 single-channel currents show a complex temporal structure, <u>as mentioned before. Moreover,</u> the open state is rather unstable, showing many rapid closures or flickers and an increased open channel noise. We observed that current flicker is more noticeable with increasing ion size and voltage (Figure 2A), suggesting that this flickery gating is a phenomenon that arises from the interaction of the permeant ion with the pore of the channel.

Increased open channel noise can occur when channels undergo very rapid transitions to closed states adjacent to the open state (Rauh et al., 2018). These dynamics can be seen as excess noise in the power spectrum of the open channel current or as a non-Gaussian distribution of the all-points current amplitude histogram (Heinemann and Sigworth, 1991a). It has been shown that for a two-state process, such as a unimolecular fast open channel block, the kinetics of the process can be characterized from all-points current amplitude histograms by means of fitting a beta distribution, which gives the amplitude of open events derived from a filtered two-state stochastic process (Fitzhugh, 1983; Yellen, 1984).

When the fast process involves more than two states, there is no exact analytic solution; however, it has been shown that the amplitude histograms can still be compared with theoretical distributions derived from a simulation of an infinite bandwidth process filtered with the same filter and frequency of experimental data. In this way, different explicit Markov models can be used to extract rate constants from extended beta distributions (Rauh et al., 2017; Schroeder, 2015). <u>Importantly, this approach is able to estimate kinetic events that are faster than the filter rise time</u> (Heinemann and Sigworth, 1991b).

Bursts of openings of TRPV1 contain both excess noise and apparently incomplete closures (Figure 5A), these two phenomena <u>make the amplitude histograms deviate from a Gaussian distribution and</u> give rise to “tails” of excess low amplitude events in the all-points current amplitude histograms. Interestingly these tails become more prominent when channels conduct larger ions.

**Figure 5.**
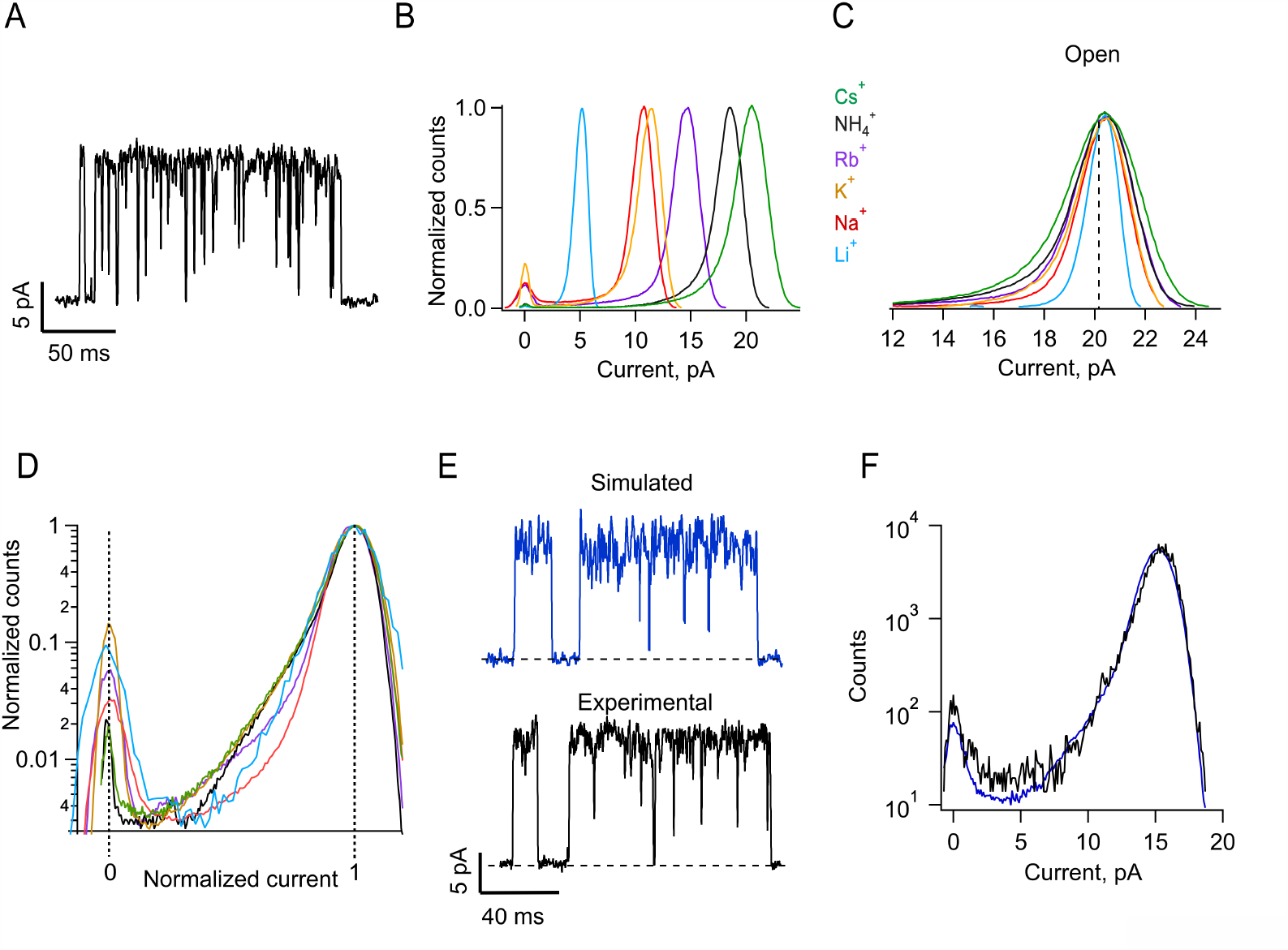
Fluctuations in the single-channel current during the open state can be fitted by a Markov model. **A)** Representative current trace during a typical single TRPV1 opening burst recorded in the inside-out configuration at 90 mV, with internal 130 mM RbCl + 100 nM capsaicin. **B)** Normalized all-points histograms showing single-channel current amplitudes of openings elicited by 100 nM capsaicin at 90 mV, for each ion tested. The histograms are shown aligned to the closed state amplitude peak. **C)** All-points histograms aligned to the maximum current amplitude of the open state to highlight the asymmetric form of the histograms. **<u>D)</u>**<u>The same histograms in C are shown normalized both in number of counts and current amplitude. This representation emphasizes the ion-dependence of the open current asymmetry.</u> **E)** Current trace of a single-channel at 90 mV from an inside-out patch exposed to internal 130 mM NH_4_Cl + 100 nM capsaicin (black) and simulation with the model in Scheme I (blue). These simulated recordings are used to produce simulated all-points amplitude histograms of single-channel current, which are then fitted to experimental all-points amplitude histograms. **F)** Typical fit of a simulated histogram (blue) to experimental histogram elicited by 110 mV with internal 130 mM NH_4_Cl + 100 nM capsaicin (black) using the Markov model in Scheme I. Parameters obtained from the fit are: α_o_ = 1500 s^-1^; α_c_ = 27 s^-1^; β_o_ = 22500 s^-1^; β_c_ = 600 s^-1^; γ_o_ = 127500 s^-1^; γ_c_ = 30000 s^-1^; χ^2^ = 0.00013873.

Figure 5B compares histograms for all ions tested at 90 mV. It can be observed that the smaller ions (Li^+^ and Na^+^) produce amplitude histograms that are roughly Gaussian, while for the larger ions, the histograms deviate from Gaussian and are asymmetric, with increased counts on the left arm of the <u>histogram contributed by amplitude values that approach the closed amplitude (Supplementary Figure 2).</u> This is made more evident in Figures 5C and D, which scale the histograms to the maximum open amplitude for all ions <u>and between the closed and open amplitudes, respectively. These figures show greater excess noise for the openings with larger cations.</u>

To extract rate constants for the fast processes that give rise to these tails, we employed extended beta distributions derived from the following Scheme I:

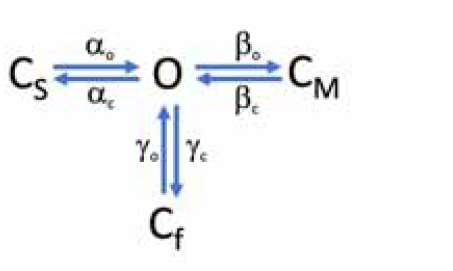

This scheme postulates the existence of three closed transitions adjacent to an open state, each transition representing a slow, medium and fast equilibrium given by alpha, beta and gamma rate constants, respectively. Although it is known that TRPV1 channels have many open and closed states, this simplified scheme can be applied to the study of bursts of openings at sub saturating capsaicin concentrations, since the effect of capsaicin is mainly to reduce the inter burst interval (Hui et al., 2003).

This scheme generates gating behavior that is very similar to the experimentally observed currents (<u>Figure 5E</u>) and the derived all-points histograms provide a good fit to the experimental histograms (<u>Figure 5F</u>). We also tested two simpler schemes with one open and one or two closed states and show that these cannot provide an adequate fit to the data (Supplementary figure 3; green and blue traces, respectively). The voltage-dependence of the forward and backward rate constants obtained with the extended beta distribution method for all ionic conditions is shown in Figure 6. The slow opening transition (α_o_) is determined mainly by longer lived closed states and shows a slight voltage-dependence in all ions, except ammonium, <u>which tends to have a higher valence</u>. Inversely, the closing transition (α_c_) shows a higher voltage-dependence. Intermediate opening and closing transitions (β_o_ and β_c_) have less voltage-dependence, while the fast transitions (γ_o_ and γ_c_) are again more voltage dependent. The voltage-dependence of these transition rate constants was fit to a single exponential function of voltage (Eq. 5). the fast-opening transitions show saturation at high positive voltages and can only be fitted with the subtraction of two exponential functions (Eq. 6), (<u>see Table II for all fitted parameter</u>). An interesting finding derived from these measurements is that the rate constants <u>for all three transitions</u> are modulated by the permeant ion. <u>When plotting the values of the rate constant at 0 mV (Figure 6B), the fast transition (γ_o_) becomes faster for openings in the presence of larger ions, although the change is not very large. In contrast, the intermediate opening rate (β_o_) clearly increase with the radius of the ion. The slow opening transition (α_o_) shows a dependence on the ionic radius that is rather interesting, being fast for small and large ions and with a minimum for intermediate size ions.</u> This slow transition mirrors the dependence of the open probability on the ionic radius (Figure 3B). In general, rate constants, when compared at 0 mV, are faster with the larger ions such as Cs^+^ and slower with smaller ions, although this is not a rule.

**Table 2.** Parameters of the fits to equation 5 of the data in Figure 6.

| Ion | Rate constants | Rate constant at 0 mV ($s^{-1}$) | $Z_1(e_0)$ |
| --- | --- | --- | --- |
| $Li^+$ | $\alpha_o$ | 1422.3 | 0.02548 |
| | $\alpha_c$ | 198.6 | -0.36236 |
| | $\beta_o$ | 2042.7 | 0.4883 |
| | $\beta_c$ | 60.337 | 0.6112 |
| | $\gamma_o$ | 87664.2 | 0.028629 |
| | $\gamma_c$ | 34744 | -0.051124 |
| $Na^+$ | $\alpha_o$ | 875.09 | -0.13178 |
| | $\alpha_c$ | 163.28 | -0.55262 |
| | $\beta_o$ | 6302.1 | 0.15402 |
| | $\beta_c$ | 191.277 | 0.30446 |
| | $\gamma_o$ | 83222 | 0.021433 |
| | $\gamma_c$ | 28800.5 | -0.02206 |
| $K^+$ | $\alpha_o$ | 291.38 | -0.20251 |
| | $\alpha_c$ | 357.04 | -0.5549 |
| | $\beta_o$ | 1000.6 | 0.312546 |
| | $\beta_c$ | 22.5 | 0.4867 |
| | $\gamma_o$ | 62298.3 | 0.10147 |
| | $\gamma_c$ | 28322 | -0.01003 |
| $Rb^+$ | $\alpha_o$ | 123.32 | -0.23134 |
| | $\alpha_c$ | 213.72 | -0.44403 |
| | $\beta_o$ | 8294.0 | 0.042546 |
| | $\beta_c$ | 210.2 | 0.24134 |
| | $\gamma_o$ | 78027.2 | 0.044316 |
| | $\gamma_c$ | 36391 | -0.11627 |
| $NH_4^+$ | $\alpha_o$ | 236.23 | 0.31361 |
| | $\alpha_c$ | 596.9 | -0.50654 |
| | $\beta_o$ | 3077.9 | 0.32154 |
| | $\beta_c$ | 33.656 | 0.582 |
| | $\gamma_o$ | 72723.3 | 0.08284 |
| | $\gamma_c$ | 24770.1 | -0.034484 |
| $Cs^+$ | $\alpha_o$ | 1543.1 | 0.014812 |
| | $\alpha_c$ | 231.63 | -0.4304 |
| | $\beta_o$ | 21545.9 | -0.055021 |
| | $\beta_c$ | 521.0 | 0.21361 |
| | $\gamma_o$ | 97846 | 0.037125 |
| | $\gamma_c$ | 34519 | 0.011 |

**Figure 6.**
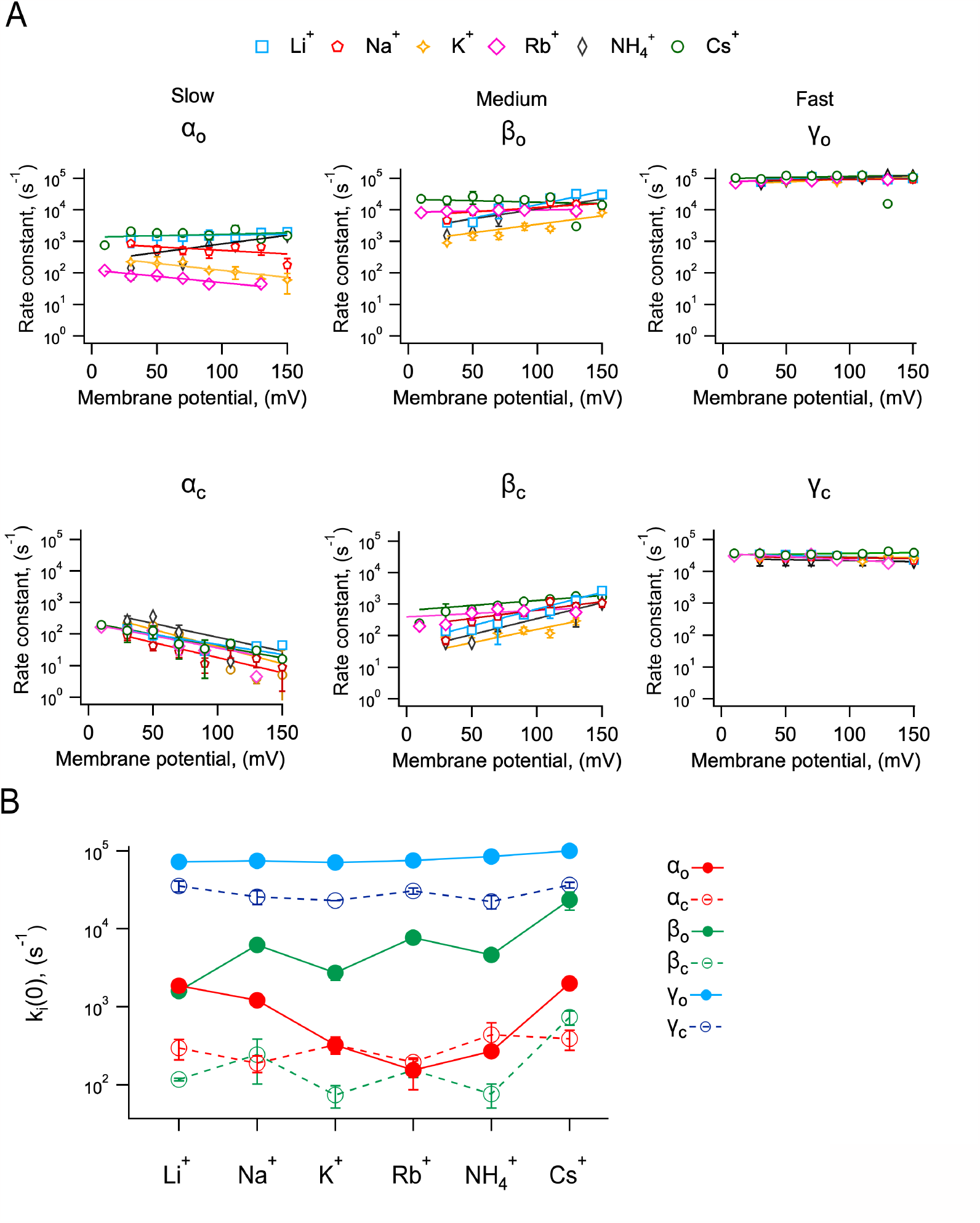
Voltage-dependent transitions near the open state of TRPV1 channels are modulated by the permeant ions. **A)** Rate constants obtained from the Markov model fit to the histograms as in Figure 5F. The rate constants were classified as slow, medium, and fast according to their magnitude at 0 mV. The continuous curves represent the fit to Eq. 5. **B)** <u>Plot of the value of the rate constants at zero mV obtained from the fit of equation 5</u> to rate constants in A. Data are mean ± s.e.m. n is: 3 (Li^+^); 6 (Na^+^); 2(K^+^); 3 (Rb^+^); 3 (NH_4_^+^) and 3 (Cs^+^).

The interplay between voltage and the effect of the permeant cation can be quantified by the total charge associated with each of the three transitions is given by: 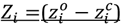, where the *z*_*i*_’s are the partial charges associated with each rate constant and *i* = α, β, γ and *o* and *c* represent the opening or closing rate constant, respectively. <u>There is no difference between the values in the different permeant ion conditions (Data not shown).</u> It is readily apparent that the smaller ions Li^+^ and Na^+^ are associated with a larger Z than say, Cs^+^ or Rb^+^.

The finding that these transitions, which occur near the open state, are voltage dependent with a relatively high associated valence, that depends on the size of the ion suggest that the observed voltage-dependence of the macroscopic conductance might arise in part from the movement of ions in the pore’s electric field. This is consistent with our previous observation that the macroscopic voltage-dependence at a saturating capsaicin concentration is modulated by the permeant ion (Figure 1D), with a tendency for smaller apparent charge for larger ions. To compare the observed voltage-dependence of rate constants with the macroscopic conductance, we measured conductance at the same concentration of capsaicin at which rate constants were estimated (100 nM) <u>from single-channel recordings</u> and contrasted it to the prediction of Scheme I, using the measured rates and charges.

The open probability predicted by Scheme I is given by <u>(derivation in Appendix):</u>

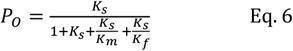

With equilibrium constants *K*_*i*_ given by:

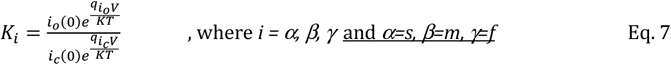

Figures 7A and B depict macroscopic currents measured at saturating (10 μM) and subsaturating (100 nM) capsaicin. Figure 7C shows G-V curves at 100 nM capsaicin for all ions and Figure 7D shows the prediction of Scheme I (Eq. 6 and 7), using values derived from measurements in Figure 6 (Table I). <u>For all ions except ammonium</u>, the model shows good agreement with the data and suggest that the estimated apparent charge associated with channel activation is modestly modulated by the permeant ion (Figure 7E).

**Figure 7.**
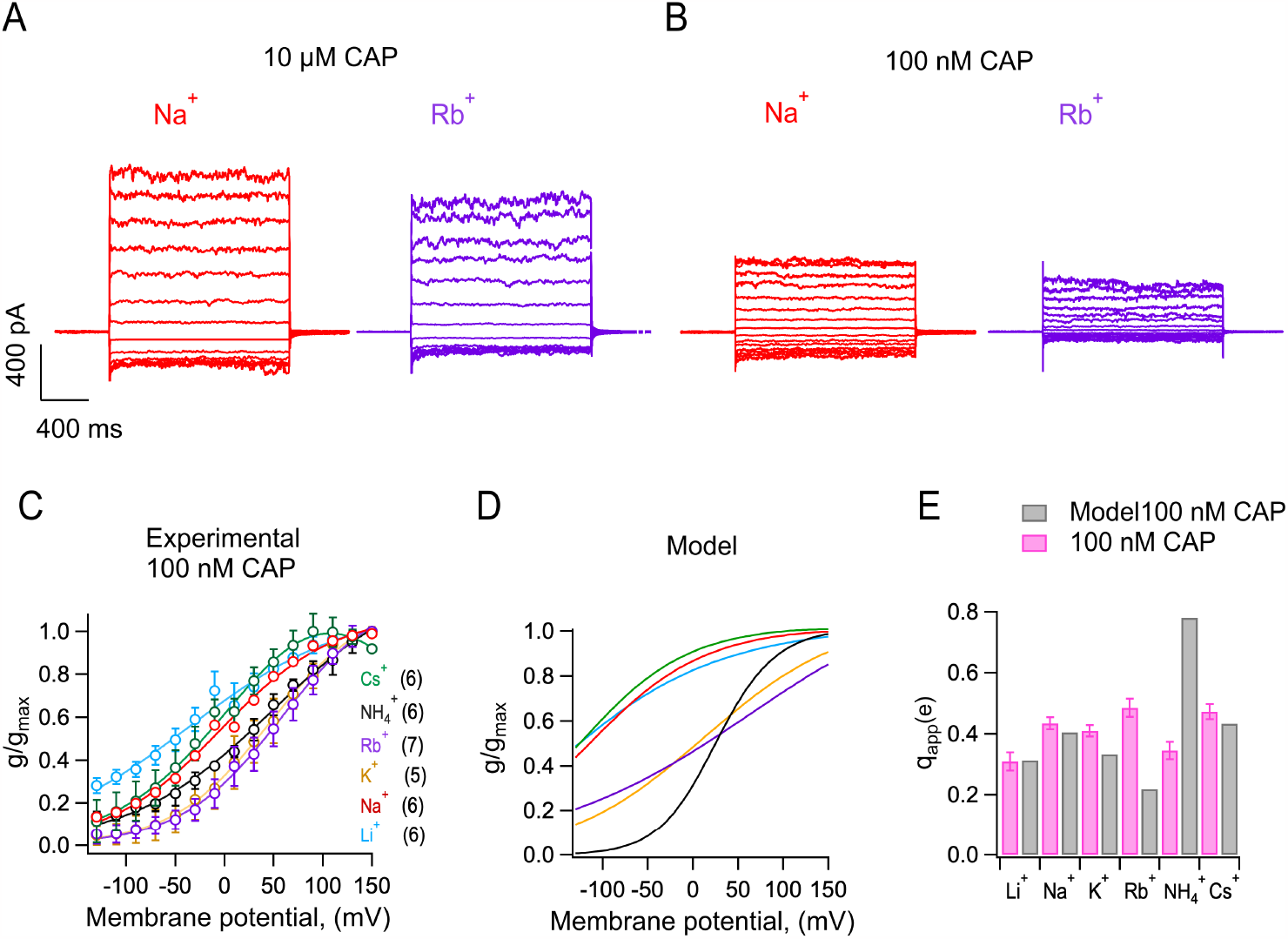
The opening probability predicted by the Markov <u>partially</u> model explains the macroscopic experimental behavior. **A)** Representative inside-out macroscopic current traces recorded in the same patch, elicited by 20 mV steps from −130 mV to +150 mV for 150 ms. The patch was exposed to internal 130 mM NaCl + 10 μM capsaicin (red) followed by 130 mM RbCl + 10 μM capsaicin (purple). **B)** Representative inside-out current traces like A but using 100 nM capsaicin. **C)** Normalized conductance-voltage (G-V) relations with monovalent ions, obtained from inside-out patches after treatment with internal 100 nM capsaicin. The G-V relation was fitted to the Boltzmann function (line). Data are shown as mean ± s.e.m., number of experiments n, in parenthesis. **D)** Normalized prediction of open probability by a Markov model for each monovalent ion. **E)** Bar graph of apparent charge (q_app_)associated with channel opening estimated by fitting a Boltzmann equation to curves in A and B. Data is shown as mean ± s.e.m.

## Discussion

TRPV1 is generally characterized as a cation permeable, non-selective channel with slightly higher permeability to Ca^2+^ ions (Caterina et al., 1997b). Surprisingly, little is known of the permeation mechanisms for cations in TRPV1. A study by (Samways and Egan, 2011) showed single-channel recordings at a negative voltages that suggested non-selective permeability of cations (also based in single-channel conductance) in the order: Na^+^ > Cs^+^ > Li^+^ > K^+^. In this study, we have employed single-channel recordings under bi-ionic conditions and measured a permeability sequence based on the conductance at positive voltages, where the single-channel conductance is higher and easier to measure. We observed a sequence Cs^+^ > Rb^+^ > K^+^ > Na^+^ > Li^+^ <u>for alkali cations</u>. Interestingly, this is identical with an Eisenman sequence of type I (Eisenman, 1962), which is produced by a weak field-strength cation binding site that favors permeation of easily dehydrated larger ions. Molecular dynamics (MD) simulations show that the TRPV1 pore may contain up to four binding sites for monovalent cation permeation in which the free energy for Na^+^ and K^+^ is very similar (Jorgensen et al., 2016), consistent with the almost identical permeability ratio for Na^+^ and K^+^. However, no definitive permeation mechanism has been dilucidated for TRPV channels.

Our data suggest that while non-selective, the pore of TRPV1 channels can regulate the flux of monovalent cations based on the energetics of dehydration and can differentiate between the size of ions, which is borne by the permeability sequence obtained from single-channel records (Figure 2C).

MD simulations support a highly flexible selectivity filter in TRPV1 (Darré et al., 2015) and our results indicated that gating can also be dynamic and is modulated by the permeant ion. These results prompted us to explore the <u>microscopic gating kinetics and its possible dependence on the permeant ion</u>. <u>We observe that single-channel kinetics are indeed modulated by the permeant ion, with Rb^+^ reducing the burst length and the open probability of the channels. The smaller ions Li^+^ and Na^+^ make it more probable for the channel to open in long duration bursts and Cs^+^ has an intermediate effect. On top of these effects on closed and open dwell times, we observed that the noise while the channel is open also varied with the permeant ion.</u>

To study in detail this behavior, we employ the extended beta distribution method to extract rate constants that arise from <u>all closure events, including fast closures that appear incomplete due to filtering,</u> since simpler detection methods such as half-amplitude threshold crossing are not appropriate to detect the contribution of rapid incomplete closings.

In single-channel recording experiments, <u>the intermediate transitions near the open state</u> seem to speed up when large ions permeate, leading to increased open channel noise. This observation could be explained if small ions like Li^+^ and Na^+^ are able to remain bound to the selectivity filter for longer times, due to their stronger electrostatic electric field, and thus stabilize the open state, reducing fluctuations. <u>In contrast, the slow transitions are faster for small and big ions, with a minimum for intermedia radii ions. This behavior is also observed in the burst length distributions estimated from half-amplitude threshold crossing. Interestingly, the open probability</u> shows a similar behavior, with a minimum for Rb^+^ ions. This is to be expected since <u>the slow transitions will contribute longer dwell times (in this case, longer closed times) and have a larger contribution to the measured Po (Fig. 3).</u>

The rapid fluctuations we observe are consistent with the existence of transitions from the open state to closed states that are adjacent to the open state. It is likely that these represent rapid pore dynamics derived from structural transitions. Cryo-EM experiments have revealed that many TRPV channels, including TRPV1, can exist in different pore conformations, even in the presence of activators (Zhang et al., 2021). Functional data have also provided evidence that TRPV channels can adopt different open conformations in the presence of different ligands (Benítez-Angeles et al., 2023; Canul-Sánchez et al., 2018).

Experiments employing single-molecule FRET in Kir channels have directly shown dynamic changes of the structure of the selectivity filter as a function of the presence of Na^+^ or K^+^ ions (Wang et al., 2019). Related experiments employing solution nuclear magnetic resonance have shown similar ion-dependent dynamics of the filter of a non-selective ion channel, which has a filter structure more closely related to TRPV1’s (Lewis et al., 2021) and the intrinsic pore dynamics of TRPV1 has been demonstrated directly by measuring the fluorescence of genetically encoded non-canonical amino acids (Steinberg et al., 2017). <u>It has also been demonstrated that the selectivity filter can undergo conformational changes, although it is not a gate for small ions</u> *<u>per se</u>* <u>(Jara-Oseguera et al., 2016).</u>

An open question in TRPV channel biophysics, specially TRPV1, is the origin of the mild voltage-dependence of the macroscopic conductance. As mentioned before, the VLSD does not support a canonical function as a voltage sensor (Palovcak et al., 2015b). It has recently been proposed that a conformational change that rearranges the position of negatively charged residues at the extracellular entrance of the selectivity filter is responsible for the voltage-dependence of TRPV1 gating, which is equivalent to the displacement of ∼0.55 elementary charges per channel <u>(Yang et al., 2020).</u> We find that in the presence of saturating capsaicin, the apparent gating charge depends on the identity of the permeant ion. The smallest alkali metal, Li^+^, is associated with a steeper G-V, with an apparent gating charge of 0.65 e_o_ as well as activation at more negative voltages.

It is known that other types of ion channels obtain their voltage-dependence from the concerted movements of permeant ions in the permeation pathway (Schewe et al., 2016; Marchesi et al., 2012; Pusch et al., 1995) and that permeant blockers, also moving in the pore’s electric field, can produce highly voltage-dependent block (Spassova and Lu, 1998). For example, in chloride ClC channels, voltage-dependence is a result of the interaction of Cl^-^ ions with a protopore gate (Richard and Miller, 1990) and in Kir channels, where permeating ions non-canonically regulate the voltage-dependence of gating in these channels (Lu et al., 2001; Kurata et al., 2010). <u>CNG channels also display a small voltage-dependence at low open probabilities, although its origin has not been clarified (Benndorf et al., 1999).</u>

A fascinating observation derived from our study is that at relatively low concentration of capsaicin, were the open probability P_o_ ∼0.4, the apparent gating charge is largest for the larger permeant cations, while at saturating concentration, with P_o_ = 0.98, the apparent gating charge is largest when the smaller ion (Li^+^) permeates.

## Acknowledgments

We thank Gisela Rangel, Manuel Hernández and Elsa Evaristo for excellent technical support. This work was supported by DGAPA-PAPIIT-UNAM grant No. IN215621 to L.D.I. and CONACyT (A1-S-8760) and DGAPA-PAPIIT-UNAM (IN200423) to T.R. M. G-A is a doctoral student from Programa de Doctorado en Ciencias Bioquímicas-UNAM and was supported by a doctoral thesis scholarship from CONACyT No. 776082.

## Supplementary figure legends

**Supplementary Figure 1.**
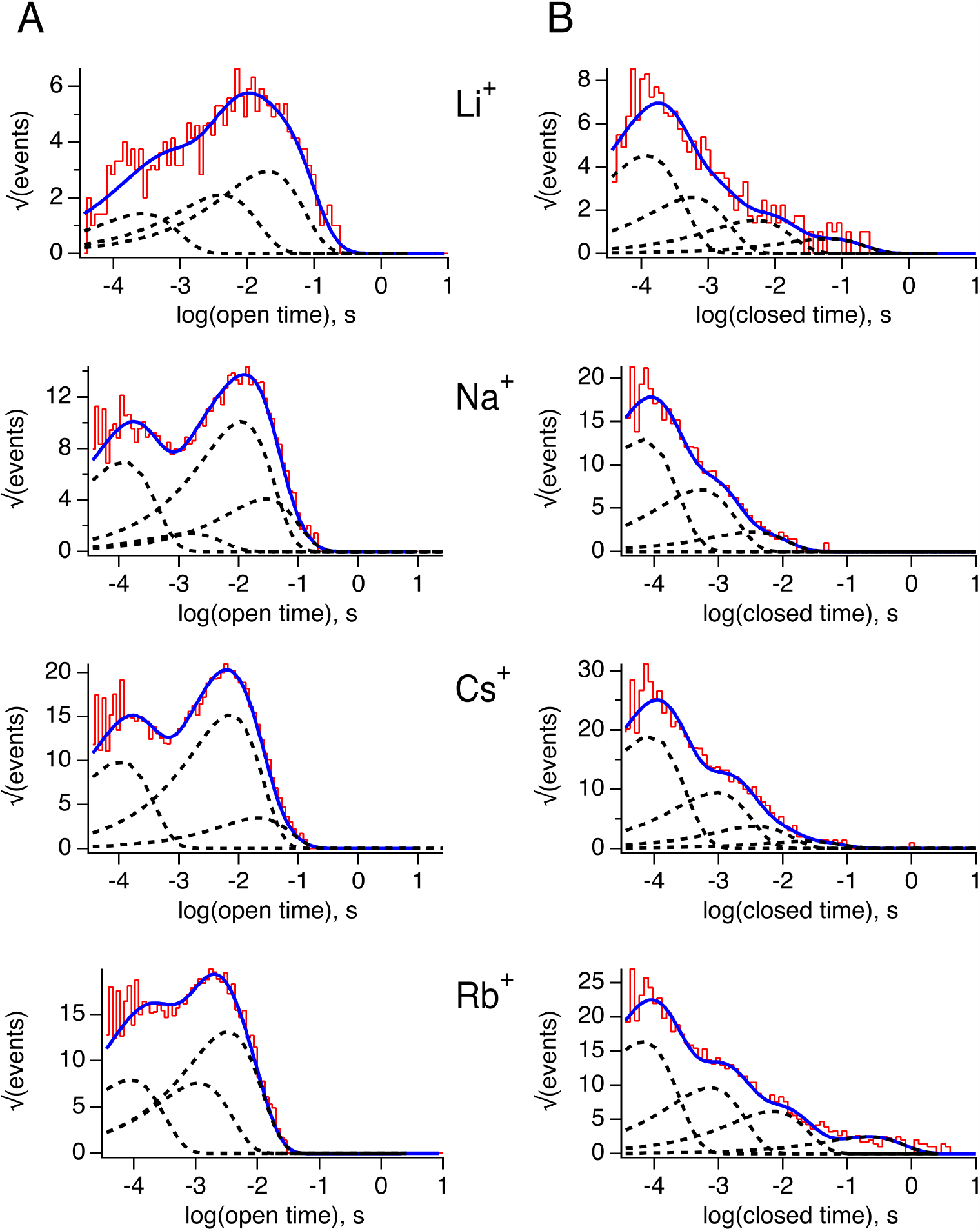
<u>Analysis of the open-closed kinetics of single-channel recordings in different permeant ions</u>. **<u>A)</u>**<u>Open time distributions in the presence of the indicated ion. The blue curve is the fit of the histogram to a sum of exponential components. Each component is indicated by the curves in dotted lines.</u> **<u>B)</u>**<u>Closed time distribution in the presence of the indicated ion. The blue curve is the fit of the histogram to a sum of exponential components. Each component is indicated by the curves in dotted lines. The values of the time constants and amplitudes of exponential components are shown in Table 1.</u>

**Supplementary Figure 2.**
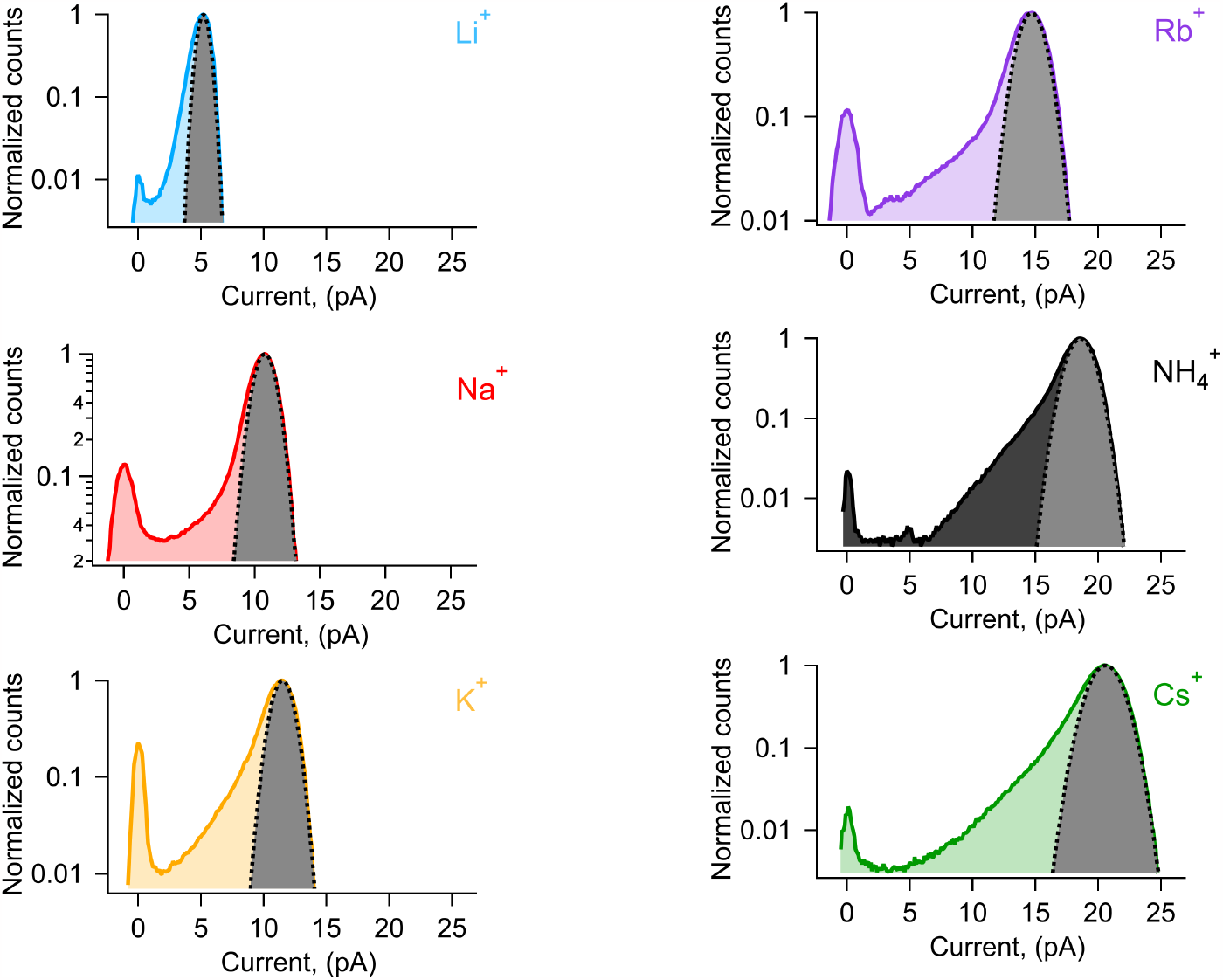
<u>Gaussian fits to amplitude histograms of currents from different ions</u>. <u>. The grey shaded and dotted curve is the best fit to each amplitude</u> histogram to a single Gaussian function. Li^+^ carried currents show the least extra open <u>channel noise, as indicated by the extra counts between the Gaussian and the peak with zero current mean that indicates the closed channel level.</u>

**Supplementary Figure 3.**
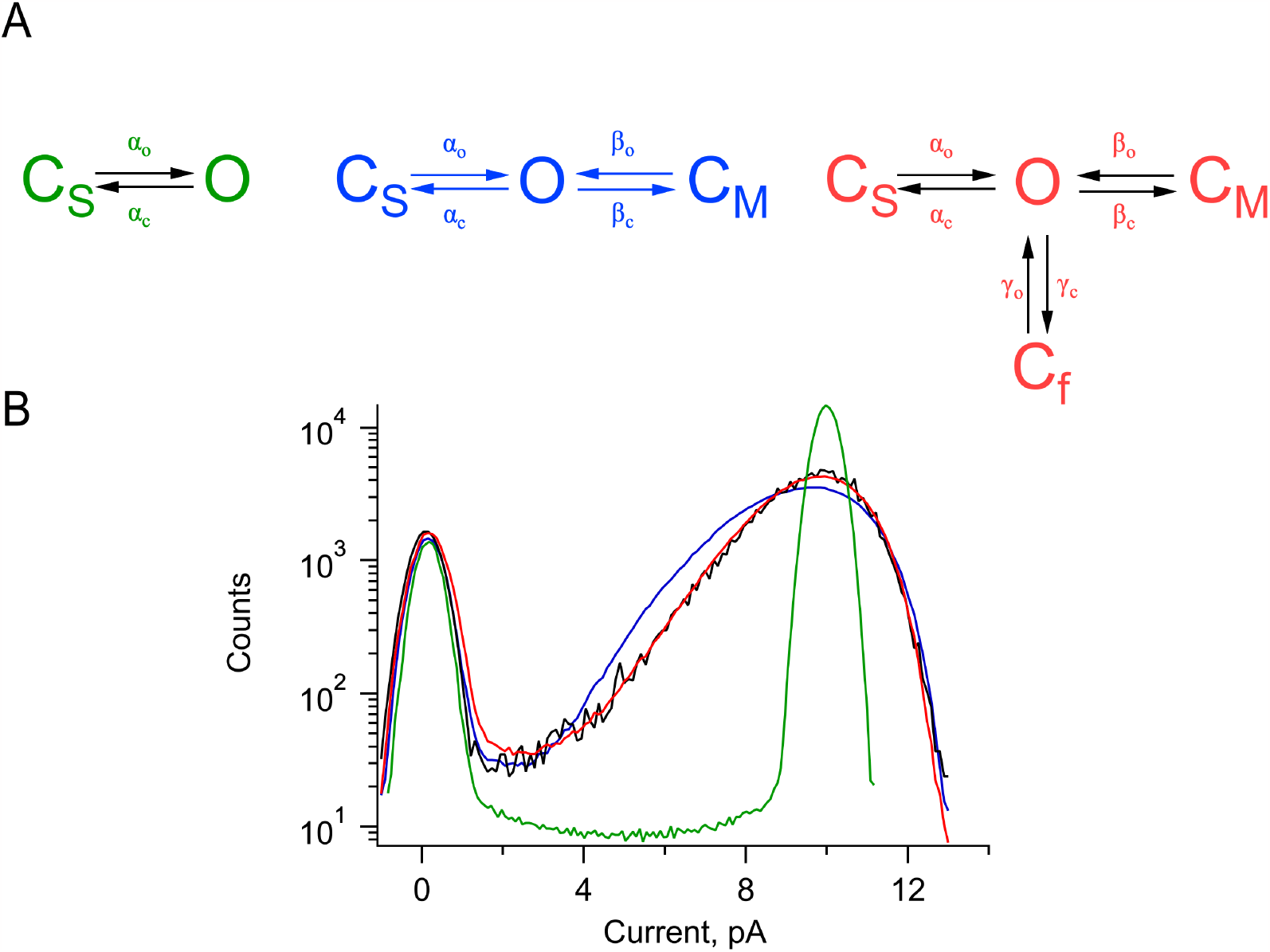
Amplitude histogram fits to three different Markov models. **A)** Markov models with one closed state (green), two closed states (blue) and three closed states (red). **B)** Fit of the simulated histograms from the models in A to experimental an all-points histogram of single channel current of TRPV1(black), elicited by a pulse to 90 mV from an inside-out patches exposed to 130 mM NaCl + 100 nM capsaicin. The fits parameters used for one close state were: α_o_ = 301.9535 s^-1^; α_c_ = 36.1664 s^-1^; χ^2^ = 0.0697; two closed states: α_o_ = 300 s^-1^; α_c_ = 37.5 s^-1^; γ_o_ = 28500 s^-1^; γ_c_ = 13500 s^-1^; χ^2^ = 0.002342; three closed states: α_o_ = 280.61 s^-1^; α_c_ = 49 s^-1^; β_o_ = 14724.475 s^-1^; β_c_ = 1133.16 s^-1^; γ_o_ = 52500 s^-1^; γ_c_ = 15396 s^-1^; χ^2^ = 0.00017296.

## Appendix: Derivation of equation 6

The equations describing the occupancy of each state in Scheme I are given by

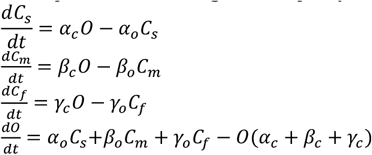

These equations are subject to: *C*_*s*_ *+ C*_*m*_ *+ C*_*f*_ *+ O = 1* and the probability of being in the open state, *P*_*o*_ is:

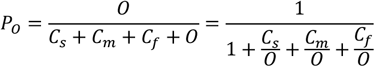

In steady-state:

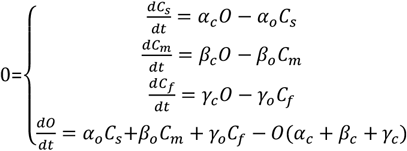

Therefore: 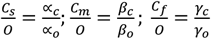 and 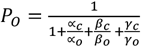

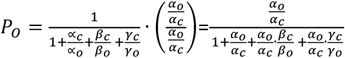. Defining 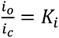 *for i* =∝, *β or γ*, we can

write:

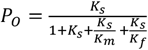, where α=*s* stands for slow, β= *m* for medium and γ= *f* for fast.

